# Quantitative analysis of HDAC inhibition using the TGx-HDACi transcriptomic biomarker in human TK6 cells

**DOI:** 10.64898/2026.11.13.766538

**Authors:** Eunnara Cho, Andrew Williams, Carole L. Yauk

## Abstract

Transcriptomic biomarkers provide an objective approach for extracting mechanistic information from complex gene expression datasets and may support quantitative applications in chemical assessment. TGx-HDACi is an 81-gene transcriptomic biomarker designed to identify histone deacetylase inhibitors (HDACi) in human TK6 cells. We evaluated whether benchmark concentrations (BMCs) derived from TGx-HDACi responses are concordant with BMCs based on reductions in HDAC enzyme activity. Human TK6 cells were exposed to concentration ranges of three HDAC inhibitors from different representative classes: the pan-HDAC inhibitors trichostatin A (TSA) and valproic acid (VPA), and the class I-selective inhibitor entinostat. HDAC enzyme activity and TGx-HDACi responses were assessed, and concentration-response modeling was performed to derive BMCs. TGx-HDACi correctly identified all three compounds as HDAC inhibitors and showed concentration-dependent responses. For TSA and VPA, TGx-HDACi and HDAC activity BMCs had overlapping confidence intervals, indicating good agreement between potency measures. In contrast, entinostat produced higher TGx-HDACi BMCs than the enzyme assay. This reduced quantitative concordance may reflect differences in HDAC selectivity, as TGx-HDACi was developed primarily using pan-HDAC inhibitors and appears less responsive to the more limited transcriptional changes induced by class I-selective inhibitors. Overall, these results demonstrate that TGx-HDACi reliably identifies HDAC inhibition and can provide potency estimates that align with enzyme activity measures for some HDAC inhibitor classes. However, the findings highlight the importance of considering biomarker applicability domain when interpreting potency estimates across compounds with differing HDAC selectivity.

## 1. Introduction

Transcriptomic biomarkers are used to objectively and efficiently extract information from complex transcriptomic data sets for chemical toxicity assessment. These biomarkers are typically comprised of a set of genes that collectively exhibit a distinct expression pattern in response to specific biological perturbations. Similarities in the expression patterns induced by a test chemical to the biomarker suggests that the chemical operates through a similar molecular initiating event or cause a similar type of toxicity (Corton et al., 2019; Krewski et al., 2020). Thus, biomarker-guided transcriptomic analyses provide valuable mechanistic information for applications in weight of evidence assessment of potential modes of action (MOAs) or hazards, or for chemical screening, prioritization, and identification of follow-up assays to streamline toxicity assessment strategies.

The TGx-HDACi transcriptomic biomarker is comprised of 81 genes that are transcriptionally altered by exposure to chemicals that cause histone deacetylase inhibition (HDACi) in TK6 human lymphoblastoid cells (Cho et al., 2021). One of the well-known effects of HDACi is hyperacetylation of histones, which alters chromatin configuration and, consequently, gene transcription. HDACs are also involved in the post-translational modification of numerous non-histone proteins that are critical to different cellular processes including RNA splicing, cell cycle, and DNA repair, as well as tissue-specific functions (Wang et al., 2020; Choudhary et al., 2009; Wright and Menick, 2016; Rahhal and Seto, 2019). The TGx-HDACi biomarker captures the early transcriptomic changes that correspond to disturbance in the acetylation status of histones and other proteins. While HDACi are used in medicine to treat primarily neurological conditions, many are teratogens that can have devastating effects on human development; therefore, unintentional exposure to HDACi in humans must be prevented (Gallinari et al., 2007; Koch et al., 2021).

The 18 known human HDACs (also referred to as lysine deacetylases or KDAC) are divided into two major categories: zinc^2+^-dependent, classical HDACs (classes I, II, and IV) and NAD^+^-dependent sirtuins, which constitute class III (Porter and Christianson, 2019). Due to the structural differences in the catalytic site, compounds that inhibit the classical HDACs do not typically affect sirtuins and vice versa. TGx-HDACi was specifically developed to detect the inhibition of the classical HDACs in classes I, II, and IV; its performance in identifying sirtuin inhibitors has not been evaluated.

Thus far, the TGx-HDACi biomarker has shown robust performance in classifying compounds as HDACi or non-HDACi in TK6 cells. TGx-HDACi correctly classified all 20 reference compounds and 11 external validation compounds that encompassed four HDACi and seven non-HDACi compounds (100% accuracy). The specificity of the biomarker extends to distinguishing HDACi from not only non-histone modulating and non-HDACi compounds, but also from other types of histone modulators such as the inhibitors of histone acetyltransferases, methyltransferases, and demethylases (Cho et al., 2021). The utility of TGx-HDACi for chemical screening and its performance in other cell lines and tissue types have indicated its potentials as a tool for discovering chemicals with previously unknown HDACi properties (Corton et al., 2022).

Transcriptomic biomarkers identify early molecular changes or molecular initiating events associated with adverse outcome pathways (AOP) (El-Masri et al., 2022). Thus far, there are three AOPs in the AOPWiki (https://aopwiki.org) that are initiated by HDACi: one of these is endorsed by the Organisation for Economic Co-operation and Development (OECD) and leads to testicular atrophy (https://aopwiki.org/aops/212) (Tanabe et al., 2021); two others that are in development lead to impairments in craniofacial development and neural tube defects. Aligning transcriptomic biomarkers to AOPs can inform the selection of appropriate follow-up assays. An important consideration is defining the quantitative relationships between key events in AOPs, which is necessary for predictive toxicology. Thus, developing an understanding of the quantitative relationships between TGx-HDACi biomarker responses and inhibition of HDAC enzyme activity is important for biomarker application in AOPs.

Herein, we examined concentration-response in TGx-HDACi biomarker genes alongside HDAC enzyme activity levels in TK6 cells to further validate this biomarker and its application in AOPs, as well as to explore its utility in quantitative analyses of HDACi compounds. We exposed TK6 cells for 4 hours to four concentrations of three prototypical HDACi compounds: trichostatin A (TSA), entinostat, and valproic acid (VPA). We measured the TGx-HDACi biomarker using TempO-Seq and HDAC enzyme activity levels using a fluorimetric assay. Both endpoints were analyzed by benchmark concentration (BMC) modeling; BMC modeling identifies the concentration at which there is a pre-defined change in response (i.e., the benchmark response, or BMR) relative to solvent controls. BMCs are used in a variety of risk assessment applications including analysis of chemical potencies (White et al., 2020). We evaluated whether BMCs derived from TGx-HDACi responses are concordant with those derived from HDAC enzyme inhibition and whether this relationship is consistent across HDAC inhibitor classes. Our work advances the utility of TGx-HDACi for quantitative analyses and potency comparison of HDAC inhibitors and help define its applicability domain.

## 2. Materials and methods

### 2.1. Chemicals

Trichostatin A (TSA) (CAS# 58,880-19-6; Active Motif Inc.) and Entinostat (Ent) (CAS# 209,783-80-2; ApexBio) were dissolved in dimethylsulfoxide (DMSO; Fisher Scientific) to prepare the treatment solutions. Valproic acid (VPA) (CAS# 1069-66-5; StressMarq BioSciences) was diluted in water.

### 2.2. Cell culture and chemical exposure

TK6 cells (ATCC# CRL-8015; ATCC, Manassas, VA) were cultured in suspension in a T75 flask with RPMI1640 medium (Gibco) supplemented with 10% v/v heat-inactivated horse serum (Gibco; New Zealand origin) and 200 µg/mL sodium pyruvate (Gibco), at 37°C in 5% CO_2_. The cell density was maintained between 1 x 10^5^ and 1 x 10^6^ cells/mL in a T75 flask.

Four concentrations of each compound were selected based on the concentration ranges that were used in the original TGx-HDACi study (Cho et al., 2021). In the 2021 study, we exposed cells to concentration ranges that spanned 8 - 10 concentrations flanking the published IC_50_ for HDAC inhibition of the three compounds (Cho et al., 2021). The expression levels of three genes that are highly responsive to HDACi (*RGN1, NEU1,* and *GPR183*) were then measured using TaqMan qPCR assays in Cho et al. (2021) and a single concentration used for biomarker development. In the present study, the first concentration that induced the maximal fold change in the three genes was set as the top concentration and an additional three concentrations within the slope of the concentration-response curve were selected (TSA: 50, 100, 200, 300 nM; VPA: 0.5, 0.75, 1, 2.5 mM; Ent: 12.5, 25, 75, 100 µM).

Prior to chemical exposure, cells were plated in either a 6-well or 96-well plate, depending on the assay. For cell viability and HDAC enzyme activity assays, cells were seeded in 96-well plates at a density of 1 x10^5^ cells/well in 99 µL of media. To generate samples for TempO-Seq, cells were plated in 6-well plates at a density of 5 x 10^5^ cells/mL in 5 mL of media. Treatment solutions were prepared in either H_2_O or dimethyl sulfoxide (DMSO) at 100x the target concentration for exposure. Treatment solutions with four different concentrations were prepared for each compound. Fifty µL or 1 µL of treatment solution or vehicle solvent were added to each well of 6-well or 96-well plates (1 % v/v) and cells were incubated at 37°C in 5% CO_2_. After 4 hours, cells in 96-well plates were removed from exposure and were either re-plated in fresh media for a 20-hour recovery period prior to cell viability assessment or were used to measure HDAC activity levels. Cells in 6-well plates were washed twice in PBS, pelleted, and stored frozen at -70 °C until RNA extraction.

### 2.3. MTT cell viability assay

The 3-(4,5-dimethylthiazol-2-yl)-2,5-diphenyltetrazolium bromide (MTT) assay (R&D Systems, Minneapolis, MN) was used to measure cell viability. Briefly, cells were exposed for 4 hours (n = 6) then allowed to recover for 20 hours in fresh medium at 37°C in 96-well plates, before light absorbance was measured on a SpectraMax microplate reader (Molecular Devices). Cell viability was calculated as the percentage of viable cells in the treated wells compared to the vehicle control wells.

### 2.4. HDAC enzyme activity assay

Relative HDAC enzyme activity levels were measured in TK6 cells after 4 hours of exposure using FLUOR DE LYS® HDAC fluorometric cellular activity assay kits (Enzo Life Sciences, Farmingdale, NY). This assay measures HDAC enzyme activity levels by the amount of the fluorescent product generated from the deacetylation of FLUOR DE LYS substrates in the cell. Briefly, TK6 cells were plated in a 96-well plate at a density of 1 x10^5^ cells/well in 99 μL of RPMI1640 medium without phenol red (Gibco) containing the FLUOR DE LYS HDAC substrate as per the manufacturer’s protocol. The cells were treated with each compound and vehicle solvent in triplicates. After 4 hours, fluorescence of the deacetylated product generated from the reaction between HDAC and the FLUOR DE LYS enzyme substrate was measured (Ex. 360 nm, Em. 460 nm) using a SpectraMax Gemini microplate reader (Molecular Devices). HDAC enzyme activity levels were measured as arbitrary fluorescence units (AFU). A one-way analysis of variance (ANOVA) with post-hoc Dunnett’s test was conducted to determine whether the change in AFU at each treatment concentration was statistically significant (p < 0.05) compared to the solvent control.

### 2.5. RNA extraction and TempO-Seq library construction

Following the 4 hour exposures, TK6 cells were harvested for RNA extraction. Total RNA was extracted and purified using the RNeasy Mini Kit (Qiagen, Toronto, ON, Canada) following the manufacturer’s protocol. RNA sample quality and quantity was assessed using a NanoDrop ND-100 spectrophotometer (Thermo Scientific, Burlington, ON, Canada) and an Agilent 2100 Bioanalyzer or an Agilent TapeStation (Agilent Technologies, Mississauga, ON, Canada). All RNA samples had A260/280 absorbance ratios of ≥ 2.0 and RNA integrity numbers (RINs) between 7.5 and 10.

A 96-sample TempO-Seq Human Whole Transcriptome Assay kit (BioSpyder, Carlsbad, CA) was used to generate sequencing libraries using total RNA from TK6 cells following the manufacturer’s protocols (Yeakley et al., 2017). Three replicates were generated on different days for whole transcriptome analysis of each chemical. Two replicates of blank controls (water only; no RNA), human universal reference RNA, and human brain total RNA (Takara Bio) were included for quality control and assessment. RNA samples (approximately 100 ng/µL) were diluted in an equal volume of the 2x TempO-Seq Lysis buffer to a final RNA concentration of approximately 50 ng/µL. Two µL of this mixture were used (RNA input of 100 ng) in library preparation. Briefly, detection oligos (DO) were annealed to RNA, unbound DOs were removed by nuclease digestion, and bound DOs were then ligated. Ligated DOs were PCR-amplified using tagged primers to generate TempO-Seq sequencing libraries. Libraries were pooled and quantified by qPCR using a KAPA SYBR FAST Universal qPCR kit for Illumina sequencing platforms (Roche, Wilmington, MA). Pooled libraries were sequenced on an Illumina NextSeq 500 using a 75-cycle flow cell (Illumina).

### 2.6. TempO-Seq data processing

BCL files were converted to FASTQ files and demultiplexed using bcl2fastq v. 2.20.0.42. FASTQ files were processed using the “pete. star. Script_v3.0” (provided by BioSpyder with the library kits). The script uses star v.2.5 to align the reads and the qCount function from QuasR to extract the feature counts specified in a Gene Transfer Format (GTF) file from the aligned reads.

We confirmed low signal (< 0.3%) from blank controls on the TempO-Seq plate. Boxplots of total mapped reads and hierarchical clustering (dissimilarity of >0.2 with the main data as a threshold) were examined to identify outliers. All samples passed the QC filter. The median number of mapped reads for the samples in the experiment was 1.5 million; the lowest sample had 0.6 million mapped reads.

The read counts were normalized to counts per million (CPM) in R 3.4.1 (Law et al., 2014). The CPM within each chemical treatment was then normalized to its vehicle control and the three replicates averaged. A cut-off of >9 was applied to the CPM read counts. The TempO-seq count data are available through the National Center for Biotechnology Information (NCBI)’s Gene Expression Omnibus (GEO accession: GSE253392).

### 2.7. TGx-HDACi biomarker analysis

The pamr function in the R statistical environment was used (www.bioconductor.org) to perform the nearest shrunken centroid (NSC) probability analysis. Briefly, the centroids for each class were generated from the expression profiles of TGx-HDACi reference chemicals belonging to each class (HDACi and non-HDACi) by averaging the expression of each gene across each class (Tibshirani et al., 2002; Cho et al., 2021). The NSC probability analysis classifies gene expression profiles as HDACi or non-HDACi based on similarities of their centroids to that of the reference chemicals; a probability cut-off of > 0.90 was applied for assigning chemicals to the HDACi class. In addition, a principal component analysis (PCA) was performed using the prcomp in R and a scatterplot was generated to represent the results.

### 2.8. Benchmark concentration (BMC) modeling

#### 2.8.1. BMC analysis of HDAC activity using PROAST

Concentration-response in HDAC enzyme activity levels were analyzed using the exponential and Hill models on the PROAST web application (https://proastweb.rivm.nl/). BMC values for a BMR of 25% was determined for all three HDACi. Model averaging was selected for BMC derivation, in which the Hill and exponential models were averaged to establish BMCs for HDAC enzyme activity level.

#### 2.8.2. BMC analysis of TGx-HDACi biomarker using BMDExpress

BMC analysis of the TGx-HDACi biomarker genes was performed using BMDExpress v2.2 using filtering criteria used in previous studies (Phillips et al., 2019; Buick et al., 2022). Biomarker genes were analyzed and filtered using the Williams trend test retaining features with a permutation p-value < 0.05 (with 100 permutations) with linear fold changes >1.5. To derive BMC_10_ values, TGx-HDACi genes that passed the pre-filters were fit to the following models: Linear, Exponential (2, 4, and 5), 2° Polynomial, and restricted Power (power restricted to ≥1). A best fit model was selected with the lowest Akaike Information Criterion (AIC) value (lowest complexity). BMCs were filtered based on the goodness of fit (p-value > 0.1), a BMC/BMCL ratio < 20, and the BMC < the highest concentration (Buick et al., 2022). The BMC values for each gene in the biomarker that passed these filters were averaged for each compound to generate the overall TGx-HDACi BMC_10_.

A BMR of 10% change from solvent control, which is approximately one standard deviation (SD) above the control, was selected, consistent with the US Environmental Protection Agency’s recommendations for conducting BMC modeling of continuous data with an uncertain level of change corresponding to an adverse effect (US EPA, 2012).

## 3. Results and Discussion

### 3.1. Concentration-dependent reduction in HDAC activity

Cytotoxicity of the selected concentrations of TSA, VPA, and Ent was evaluated using the MTT assay after 4-hour exposures followed by a 20-hour recovery period in fresh media (Supplementary Figure 1). Under these exposure conditions, none of the compounds induced a detectable reduction in cell viability compared to the vehicle control-treated cells.

We then used the FLUOR DE LYS assay to establis levels of HDAC enzyme activity after 4 hours of exposure (Figure 1). All three HDACi compounds induced a concentration-dependent reduction in HDAC activity levels; this reduction was statistically significant at the highest concentration of both VPA and TSA, while all tested concentrations of Ent significantly decreased HDAC activity compared to the vehicle control (p < 0.05).

**Figure 1.**
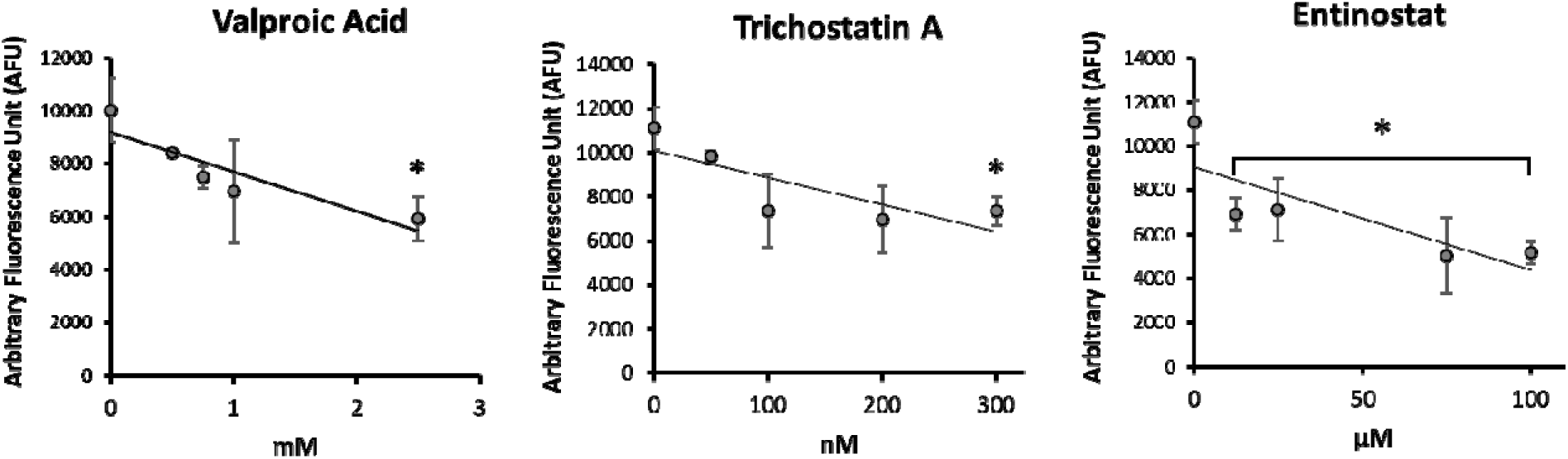
HDAC enzyme activity measured in TK6 cells after 4 hours of exposure to valproic acid (left), trichostatin A (middle), and entinostat (right) using the FLUOR DE LYS assay (n = 3). Error bars represent standard deviation. Asterisk (*) indicates statistical significance (p-value < 0.05) in one-way ANOVA with post-hoc Dunnett’s test. For entinostat, all concentrations within the indicated range caused a statistically significant change in AFU.

Ent is a selective HDACi that targets three HDACs in class I, while TSA and VPA inhibit all classical HDACs in both classes I and II (Morris and Monteggia, 2013). Despite the class-specific inhibition by Ent, it induced a reduction in the overall deacetylase activity comparable to that of the two broad spectrum HDACi in the TK6 cells (53% reduction in fluorescence at the highest concentration of Ent vs. 41% and 34% for VPA and TSA, respectively). Class I HDACs are ubiquitously expressed across different tissue types, while class II HDACs are expressed in a tissue-specific manner. In addition, class II HDACs have a comparatively lower catalytic activity than class I HDACs (Lahm et al., 2007; Bahl and Seto, 2021). It is possible that class II HDACs are the predominant HDACs in TK6 cells and perform a large portion of lysine deacetylation in these cells, since the selective inhibition of class I HDACs in TK6 cells by Ent induced a comparable level of reduction in the overall deacetylase activity as the two pan-HDACi. Overall, our data support that all three of the compounds produced concentration-dependent reductions in HDAC activity at concentrations that do not reduce cell viability.

### 3.2. TGx-HDACi biomarker analysis

We first evaluated whether TGx-HDACi classifications were consistent with the HDAC inhibition observed in the enzyme assay and whether transcriptomic profiles provided additional mechanistic insight. Each concentration of the compounds was classified as HDACi or non-HDACi using: (a) the NSC probability analysis with a 90% probability cut-off (Figure 2); or (b) a PCA with the TGx-HDACi reference compounds from Cho et al. 2021 (Figure 3).

**Figure 2.**
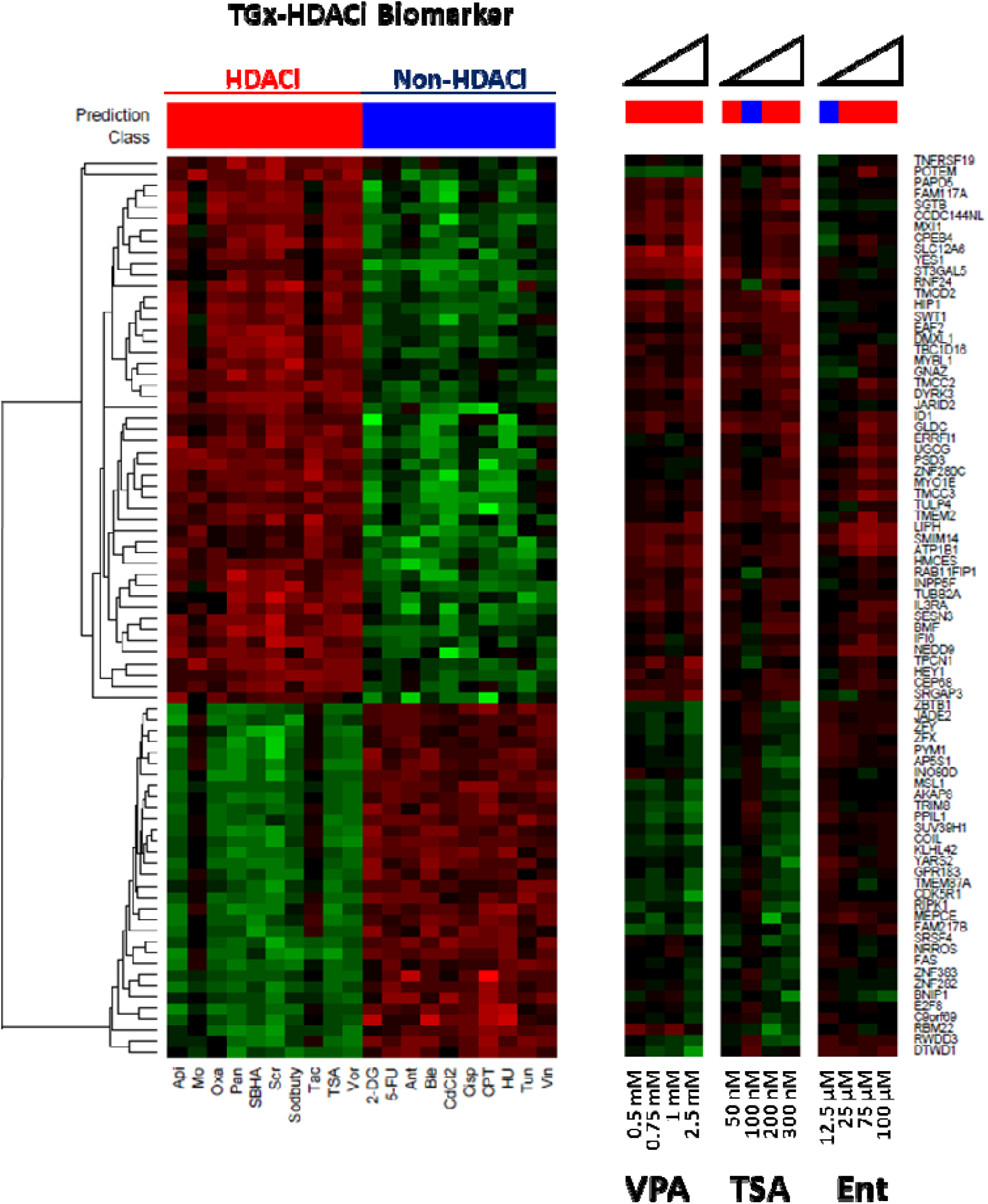
Heat maps of the TGx-HDACi biomarker (left panel) and the biomarker measured in TK6 cells after 4 hour exposures (right panels) to valproic acid (VPA; left), trichostatin A (TSA; middle), and entinostat (Ent; left) by TempO-Seq (n = 3). The 81 genes of TGx-HDACi are listed across the Y-axis and the compounds are on the X-axis. Red indicates up-regulation and green indicates down-regulation in gene expression compared to vehicle control. The bars above the heat maps show the predictions made using TGx-HDACi in a nearest shrunken centroid (NSC) probability analysis with red indicating HDACi (90% cut-off) and blue indicating non-HDACi.

**Figure 3.**
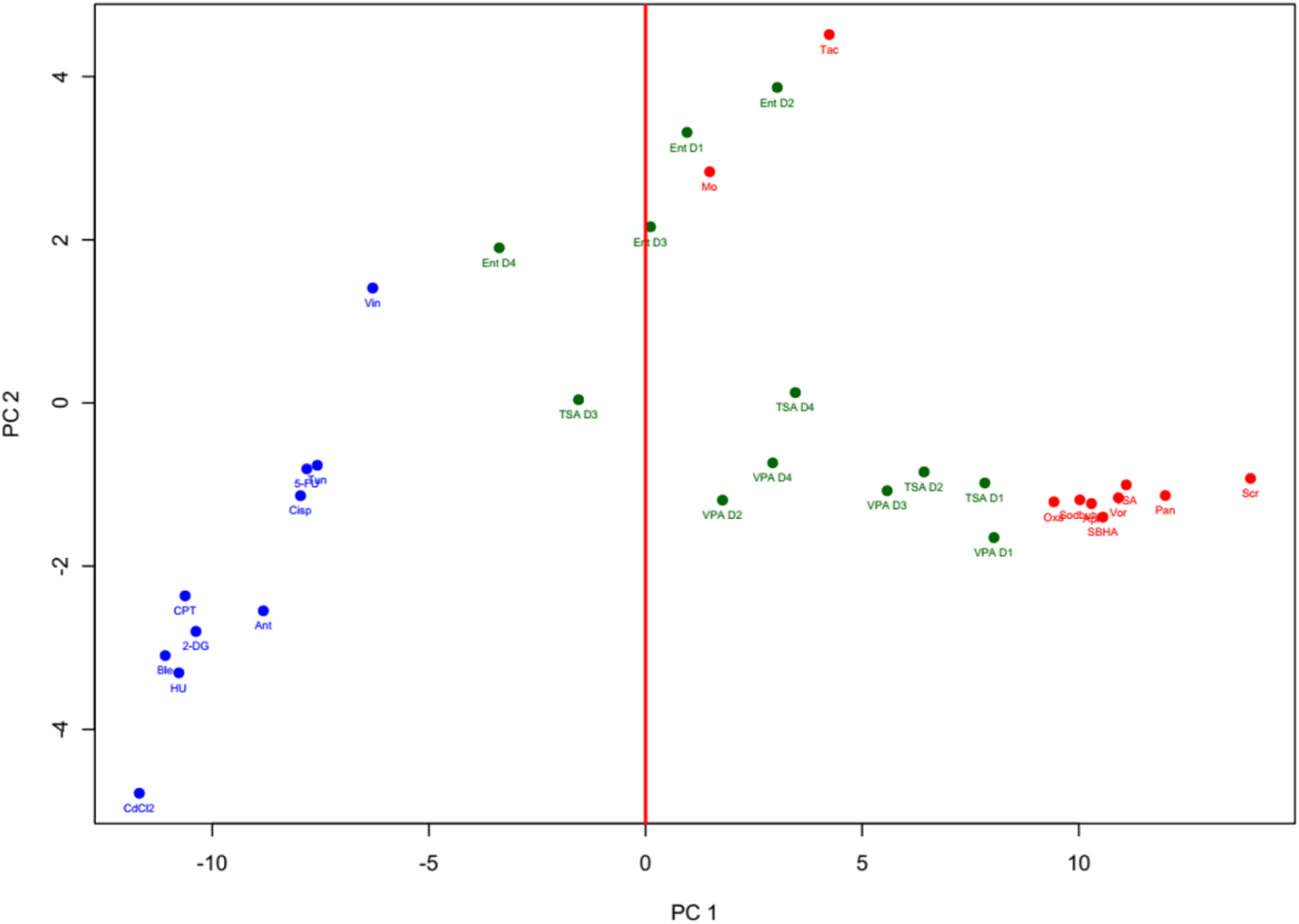
Principal component analysis of TempO-Seq gene expression profiles of the 81 genes of the TGx-HDACi biomarker measured in TK6 cells after 4 hour exposures (n = 3). Blue indicates non-HDACi reference compounds, red indicates HDACi reference compounds, and green represents the four concentrations of the three HDACi compounds used in concentration-response analysis (D1 – highest concentration; D4 – lowest concentration).

All of the concentrations of VPA produced HDACi calls in the NSC probability analysis. The second lowest concentration of TSA and the lowest concentration of Ent yielded non-HDACi calls, while all remaining concentrations of the two compounds produced HDACi classifications. The clustering of the gene expression profiles in the PCA was consistent with the NSC classifications for all three compounds; the TSA and Ent concentrations (one each) that were classified as non-HDACi in the probability analysis were on the non-HDACi side in the PCA (negative PC1) with the non-HDACi reference compounds. All concentrations of VPA clustered with the reference HDACi (positive PC1). Therefore, classification based on TGx-HDACi was more sensitive for VPA and TSA than the enzyme assay (which only showed the top concentrations as HDACi), but marginally less sensitive for Ent as it classified the lowest concentration as non-HDACi. The results support the high sensitivity of TGx-HDACi in identify concentrations at which HDACi may be occurring.

In addition to classification, the PCA revealed a distinct secondary cluster within the HDACi group. The main cluster (bottom right of Figure 3) consists of non-selective, pan-HDACi compounds, including TSA (3 concentrations) and VPA (all concentrations). In contrast, Ent clustered with the only other two class I-selective inhibitors within the TGx-HDACi reference set (Mocetinostat (Mo) and tacedinaline (Tac)) (top right Figure 3). Thus, under the exposure conditions used in this study, TGx-HDACi could distinguish the two groups of HDACi with different specificities. The results indicate that additional insights into mechanisms may be gained through analyzing transcriptomic data using different statistical approaches.

Differential gene expression patterns induced by class I-selective HDACi versus pan-HDACi have been described previously (Rajan et al., 2018). Class I HDACs are mostly present in the nucleus and class II HDACs shuttle between the cytoplasm and the nucleus, with each HDAC having a different set of deacetylation targets in the cell (Park and Kim, 2020; Bahl and Seto, 2021). Knockdown of individual HDACs has been shown to cause an upregulation or downregulation of different proteins and affect different pathways (Morris and Monteggia, 2013; Rajan et al., 2018). Consistent with these observations, TGx-HDACi captures common transcriptional responses associated with HDAC inhibition while retaining sufficient resolution to distinguish between the two HDACi inhibitor classes. These results suggest that inhibitor selectivity is an important determinant of biomarker behavior and may contribute to differences in quantitative concordance between transcriptomic and enzymatic endpoints.

### 3.3 BMC analysis of transcriptomic responses and HDAC enzyme activity

The primary objective of this study was to determine whether the concentration-response relationships in the TGx-HDACi biomarker could be modeled to derive BMCs, thereby broadening its application in AOP development/application and chemical potency analysis. We first examined the concordance between BMCs derived from TGx-HDACi responses and those derived from HDAC enzyme activity levels.

Concentration-response was first modeled for HDAC enzyme activity to derive BMCs. Although a 10% BMR is commonly used for bioassays (U.S. Environmental Protection Agency, 2012), a BMC could not be derived for Ent at this response level. Thus, we applied a 25% BMR for our analysis to enable comparison across all three compounds. Using this approach, the BMC_25_ (25% reduction in HDAC enzyme activity levels compared to control) for VPA and TSA were very close to the lowest test concentration of each compound, while the BMC_25_ for Ent was marginally (3.6-fold) below the lowest concentration used in the study (Table 1).

**Table 1.** Benchmark concentrations (BMCs) and confidence intervals (CIs) for various benchmark response levels in HDAC enzyme activity for valproic acid (VPA), trichostatin A (TSA), and entinostat (Ent).

|  | Average BMC<sub>25</sub> and 90% CI (μM)<br>(Average of Hill and Exponential models) |  |  |
|---|---|---|---|
| Compound | BMC | BMCL | BMCU |
| VPA | 737 | 287 | 1440 |
| TSA | 0.07 | 0.05 | 0.12 |
| Ent | 3.5 | 0.11 | 19.2 |

A BMR of 10% was applied to derive BMCs for the TGx-HDACi biomarker genes. Concentration-responses in individual genes were first modeled to derive gene-specific BMCs, which were then averaged to produce the composite biomarker BMC for each compound. The number of genes that could be modeled and met the filtering criteria varied by compound (Figure 4 and Table 2). Consistent with the observed potency differences, BMC_10_ could be derived for 35 of the 81 TGx-HDACi genes for TSA, the most potent of the three compounds, compared with 17 genes for Ent and 13 genes for VPA, the least potent HDACi of the three. Importantly, the relative potency ranking by the enzymatic assay and TGx-HDACi BMCs were identical (TSA > Ent > VPA) (Figure 5), indicating that TGx-HDACi captured the relative ordering of compound potency despite differences in the magnitude of the derived BMCs.

**Figure 4.**
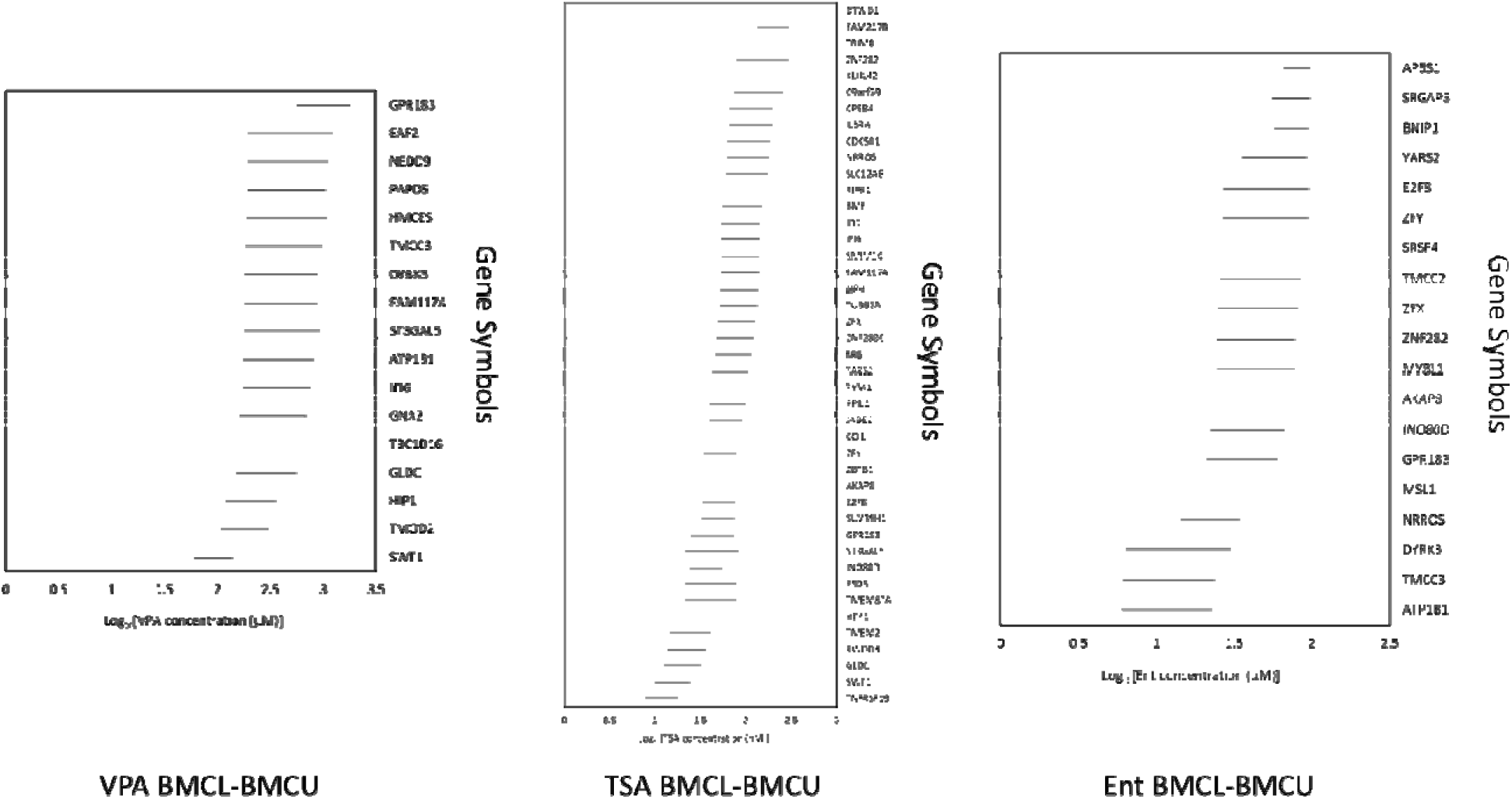
Benchmark Concentration Lower (BMCL) and BMC Upper (BMCU) confidence intervals for valproic acid (left: VPA), trichostatin A (middle: TSA), and entinostat (right: Ent) for TGx-HDACi biomarker genes that fit BMC models (gene symbols shown on y-axis) (n=3). BMDExpress software was used to derive the BMC values using a benchmark response of 10%.

**Figure 5.**
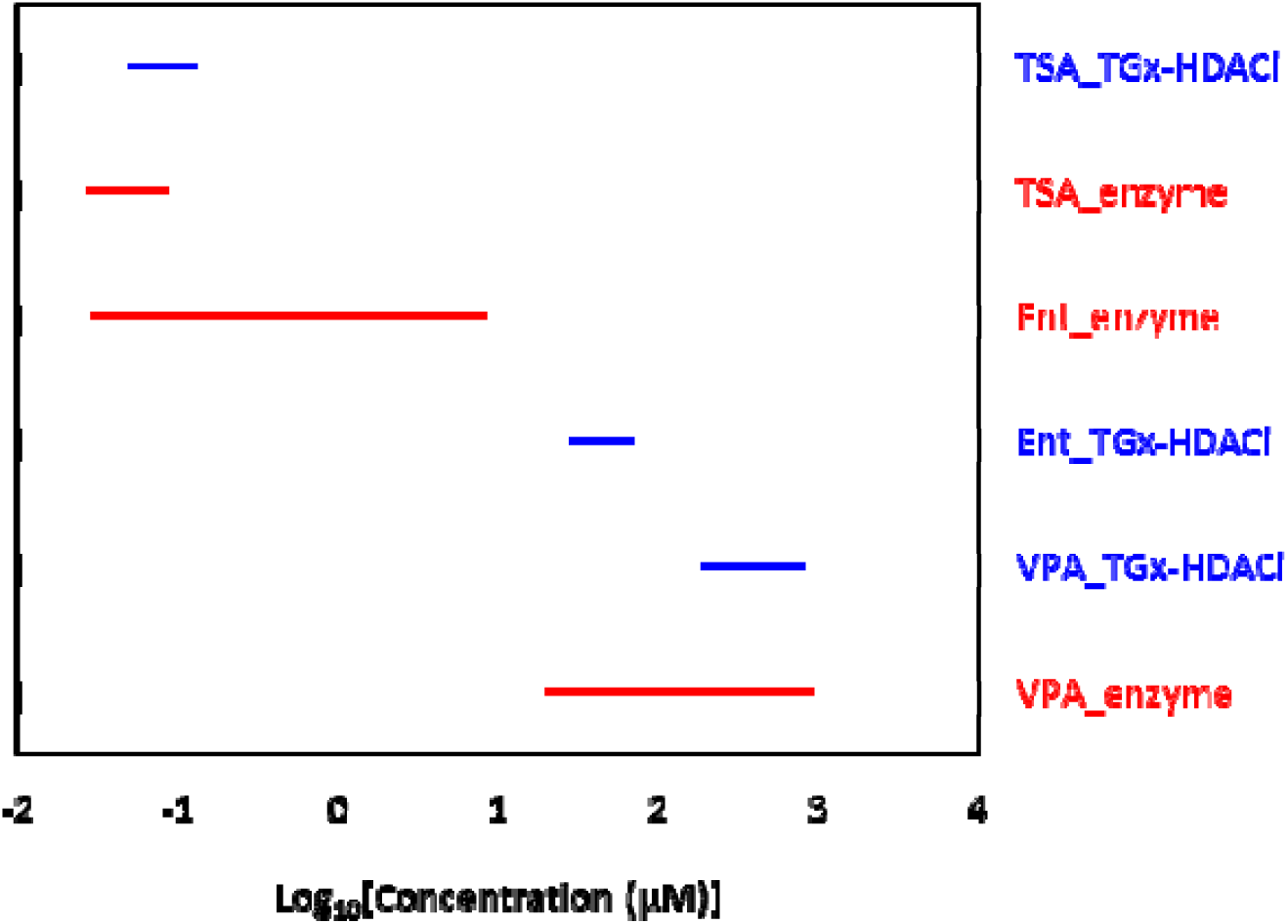
Benchmark concentrations (BMC) and 90% confidence intervals for a benchmark response of 10% (TGx-HDACi; blue) or 25% (enzyme activity; red) in TK6 cells exposed to valproic acid (VPA), trichostatin A (TSA), and entinostat (Ent) for 4 hours.

**Table 2.** Benchmark concentrations (BMC), confidence intervals (CI), and the number of TGx-HDACi genes modeled for valproic acid (VPA), trichostatin A (TSA), and entinostat (Ent).

|  | Average BMC<sub>10</sub> and 90% CI of TGx-HDACi genes (µM) |  |  |  |
|---|---|---|---|---|
| Compound | Number of genes modeled | BMC<sub>10</sub> | BMCL | BMCU |
| VPA | 13 | 125 | 53 | 374 |
| TSA | 35 | 0.069 | 0.047 | 0.12 |
| Ent | 17 | 41 | 27 | 78 |

To compare the two endpoints, the 90% confidence intervals of the BMCs for both TGx-HDACi and HDAC enzyme activity were aligned (Figure 5). The BMC_25_ for HDAC enzyme activity and BMC_10_ for TGx-HDACi of TSA and VPA were virtually identical, with highly overlapping confidence intervals. In contrast, Ent produced a significantly higher TGx-HDACi BMC confidence interval than the enzyme assay. Nevertheless, the BMCs were within one order of magnitude of each other, and the relative potency order was preserved. It is generally recommended that transcriptomic points of departure be interpreted at the order-of-magnitude level when comparing potency estimates across biological endpoints (Ewald et al., 2022). Overall, the analysis demonstrates very high quantitative concordance between transcriptomic and enzymatic measures of potency for the two pan-HDAC inhibitors, TSA and VPA, whereas reduced concordance was observed for the class I-selective HDAC inhibitor Ent. The results are nevertheless consistent with the concentration relationships expected between upstream HDAC inhibition and downstream transcriptional changes under the modified Bradford Hill criteria applied in AOP development.

Previous studies have investigated concordance in transcriptomic BMCs and related cellular phenotypic endpoints *in vitro*. For example, Buick et al. compared concentration-response to genotoxic agents using the TGx-DDI transcriptomic biomarker of DNA damage and the *in vitro* micronucleus test or the high-throughput CometChip in TK6 and HepaRG cell lines, respectively (Buick et al., 2021; Buick et al., 2022; Buick et al., 2017). These authors show that TGx-DDI BMC confidence intervals highly overlap between the assays and yield similar potency rankings. Thus, our work adds to the body of evidence that transcriptomic endpoint BMCs are predictive of the concentration at which targeted cellular effects of interest arise. Such studies are important for validating the biomarker as well as for exploring biomarker limitations and areas of applicability within the context of chemical assessment.

TGx-HDACi was derived using gene expression profiles of eight non-selective, pan-HDACi and two class I-specific HDACi. While TGx-HDACi is capable of detecting class I-specific HDACi (as shown in Cho et al. 2021) and in the TGx-HDACi analysis performed herein, the biomarker genes may generally be more responsive to pan-HDACi due to the much larger representation of pan-HDACi in the reference set. Indeed, in the heatmap of the TGx-HDACi biomarker, the class-I specific HDACi (Tac, Mo, and Ent) were distinguishable by the comparatively smaller fold changes in the biomarker genes they induced or downregulated. Selective inhibition of class I HDAC may induce stronger responses in a different set of genes not included in the TGx-HDACi panel. A reduced sensitivity toward class-selective HDACi should be noted as a potential limitation of the biomarker. Nonetheless, the TGx-HDACi genes are still able to capture a sufficient amount of the gene expression changes induced by class I HDACi to produce positive HDACi results in the biomarker assay. These findings suggest that HDAC selectivity may influence quantitative biomarker performance and should be considered when interpreting transcriptomic potency estimates.

## 4. Conclusion

We investigated whether the TGx-HDACi biomarker can be applied for quantitative studies of HDACi compounds. Our results support the use of concentration-response modeling to derive quantitative estimates of HDACi potency using TGx-HDACi. While strong concordance between transcriptomic and enzymatic potency estimates was observed for the pan-HDAC inhibitors examined, a somewhat reduced concordance for a class I-selective inhibitor suggests that HDAC selectivity should be considered when interpreting TGx-HDACi biomarker potency estimates. Under the tested exposure conditions, our results suggest that TGx-HDACi supports quantitative analyses of suspected HDAC inhibitors, including potency comparisons and applications within the AOP/MOA frameworks. Overall, this study identifies an additional utility of TGx-HDACi and highlights the importance of considering biomarker applicability domain in chemical assessment.

## Author Contributions

Conceptualization – EC, AW, CLY. Funding acquisition – CLY. Investigation – EC. Formal analysis – EC and AW. Visualization – EC and AW. Writing – original draft – EC. Writing – review and editing – EC, AW, CLY.

## Acknowledgments

This research was undertaken thanks to funding to CLY from the Health Canada Genomics R&D Initiative, the Canada Research Chairs Program (CRC-2020-00060) and the Natural Sciences and Engineering Research Council of Canada (NSERC) (RGPIN-2021-02806). EC was supported by NSERC’s CREATE REACT program. We would like to thank Remi Gagne and Dr. Lauren Bradford for their bioinformatics support and Drs. Lauren Bradford and Allison Loan (Health Canada) critical review of the manuscript with very helpful feedback.

## Abbreviations

AOP: Adverse outcome pathway
BMC: Benchmark concentration
BMR: Benchmark response
HDACi: Histone deacetylase inhibitor/inhibition
MOA: mode of action
NSC: Nearest shrunken centroid
PCA: Principal component analysis
POD: Point of departure
TempO-Seq: Templated Oligo-Sequencing

## References

Bahl S, Seto E. 2021. Regulation of histone deacetylase activities and functions by phosphorylation and its physiological relevance. Cell Mol Life Sci 78:427–445.

Buick JK, Rowan-Carroll A, Gagné R, Williams A, Chen R, Li HH, Fornace Jr AJ, Chao C, Engelward BP, Frötschl R, Ellinger-Ziegelbauer H, Pettit SD, Aubrecht J, Yauk CL. 2022. Integrated Genotoxicity Testing of three anti-infective drugs using the TGx-DDI transcriptomic biomarker and high-throughput CometChip® assay in TK6 cells. Front Toxicol 4:991590.

Buick JK, Williams A, Kuo B, Wills JW, Swartz C, Recio L, Li HH, Fornace Jr. AJ, Aubrecht J, Yauk CL. 2017. Integration of the TGx-28.65 genomic biomarker with the flow cytometry micronucleus test to assess the genotoxicity of disperse orange and 1,2,4-benzenetriol in human TK6 cells. Mutat Res Fund Mol Mech Mutagen 806:51–62.

Buick JK, Williams A, Meier MJ, Swartz CD, Recio L, Gagné R, Ferguson SS, Engelward BP, Yauk CL. 2021. A Modern Genotoxicity Testing Paradigm: Integration of the High-Throughput CometChip® and the TGx-DDI Transcriptomic Biomarker in Human HepaRG™ Cell Cultures. Front Public Health 9:694834.

Cho E, Rowan-Carroll A, Williams A, Corton JC, Li HH, Fornace Jr. AJ, Hobbs CA, Yauk CL. 2021. Development and Validation of the TGx-HDACi Transcriptomic Biomarker to Detect Histone Deacetylase Inhibitors in Human TK6 Cells. Arch Toxicol 95:1631–1645.

Choudhary C, Kumar C, Gnad F, Nielsen ML, Rehman M, Walther TC, Olsen JV, Mann M. 2009. Lysine acetylation targets protein complexes and co-regulates major cellular functions. Science 325:834–840.

Corton, J.C., Kleinstreuer, N.C., Judson, R.S., 2019. Identification of potential endocrine disrupting chemicals using gene expression biomarkers. Toxicology and Applied Pharmacology 380, 114683. 10.1016/j.taap.2019.114683

Corton JC, Liu J, Williams A, Cho E, Yauk CL. 2022. A gene expression biomarker identifies inhibitors of two classes of epigenome effectors in a human microarray compendium. Chem Biol Interact 365:110032.

El-Masri H, Friedman KP, Isaacs K, Wetmore BA. 2022. Advances in computational methods along the exposure to toxicological response paradigm. Toxicol Appl Pharmacol 450:116141.

Ewald, J.D., Basu, N., Crump, D., Boulanger, E., Head, J., 2022. Characterizing Variability and Uncertainty Associated with Transcriptomic Dose–Response Modeling. Environ. Sci. Technol. 56, 15960–15968. 10.1021/acs.est.2c04665

Gallinari P, Di Marco S, Jones P, Pallaoro M, Steinkuhler D. 2007. HDACs, histone deacetylation and gene transcription: from molecular biology to cancer therapeutics. Cell Research 17:195–211.

Koch BE, Spaink HP, Meijer AH. 2021. A quantitative in vivo assay for craniofacial developmental toxicity of histone deacetylase. Toxicol Lett 342:20–25.

Krewski, D., Andersen, M.E., Tyshenko, M.G., Krishnan, K., Hartung, T., Boekelheide, K., Wambaugh, J.F., Jones, D., Whelan, M., Thomas, R., Yauk, C., Barton-Maclaren, T., Cote, I., 2020. Toxicity testing in the 21st century: progress in the past decade and future perspectives. Arch Toxicol 94, 1–58. 10.1007/s00204-019-02613-4

Lahm A, Paolini C, Pallaoro M, Nardi M, Jones P, Neddermann P, Sambucini S, Bottomley M, Lo Surdo P, Carfı A. 07. Unraveling the hidden catalytic activity of vertebrate class IIa histone deacetylases. Proceedings of the National Academy of Sciences 104:17335–17340.

Law C, Chen Y, Shi W, Smyth G. 2014. voom: precision weights unlock linear model analysis tools for RNA-seq read counts. Genome Biol 15:R29.

Li HH, Hyduke DR, Chen R, Heard P, Yauk CL, Aubrecht J, Fornace Jr. AJ. 2015. Development of a Toxicogenomics Signature for Genotoxicity Using a Dose-Optimization and Informatics Strategy in Human Cells. Environ Mol Mutagen 56:505–519.

Morris MJ, Monteggia LM. 2013. Unique functional roles for class I and class II histone deacetylases in central nervous system development and function . Int J Dev Neurosci 31:370–381.

Park SY, Kim JS. 2020. A short guide to histone deacetylases including recent progress on class II enzymes. Exp Mol Med 52:204–212.

Phillips JR, Svoboda DL, Tandon A, Patel S, Sedykh A, Mav D, Kuo B, Yauk CL, Yang L, Thomas RS, Gift JS, Davis JA, Olszyk L, Merrick BA, Paules RS, Parham F, Saddler T, Shah RR, Auerbach SS. 2019. BMDExpress 2: enhanced transcriptomic dose-response analysis workflow. Bioinformatics 35:1780–1782.

Porter NJ, Christianson DW. 2019. Structure, Mechanism, and Inhibition of the Zinc-Dependent Histone Deacetylases. Curr Opin Struct Biol 59:9–18.

Rahhal R, Seto E. 2019. Emerging roles of histone modifications and HDACs in RNA splicing. Nucleic Acids Res 47:4911–4926.

Rajan A, Shi H, Xue B. 2018. Class I and II Histone Deacetylase Inhibitors Differentially Regulate Thermogenic Gene Expression in Brown Adipocytes. Sci Rep 8:13072.

Tanabe S, Hirose A, Yamada T. 2021. Adverse Outcome Pathway on histone deacetylase inhibition leading to testicular atrophyOECD Series on Adverse Outcome Pathways, No. 17. Paris: OECD Publishing.

Tibshirani R, Hastie T, Balasubramanian N, Chu G. 2002. Diagnosis of multiple cancer types by shrunken centroids of gene expression. Proc Natl Acad Sci USA 99:6567–6572.

U.S. Environmental Protection Agency. 2012. Benchmark Dose Technical Guidance. 100-R-12-001 edition. Washington, DC: U.S. Environmental Protection Agency.

Wang P, Wang Z, Liu J. 2020. Role of HDACs in normal and malignant hematopoiesis. Mol Cancer 19:5.

White PA, Long AS, Johnson GE. 2020. Quantitative Interpretation of Genetic Toxicity Dose-Response Data for Risk Assessment and Regulatory Decision-Making: Current Status and Emerging Priorities. Environ Mol Mutatgen 61:66–83.

Wright LH, Menick DR. 2016. A class of their own: exploring the nondeacetylase roles of class IIa HDACs in cardiovascular disease. Am J Physiol Heart Circ Physiol 311:H199–H206.

Yeakley J, Shepard P, Goyena D, VanSteenhouse H, McComb J, Seligmann B. 2017. A trichostatin A expression signature identified by TempO-Seq targeted whole transcriptome profiling. PLoS One 12:e0178302.

