## Supplementary Figure 1 for "Quantitative analysis of HDAC inhibition using the TGx-HDACi transcriptomic biomarker in human TK6 cells"

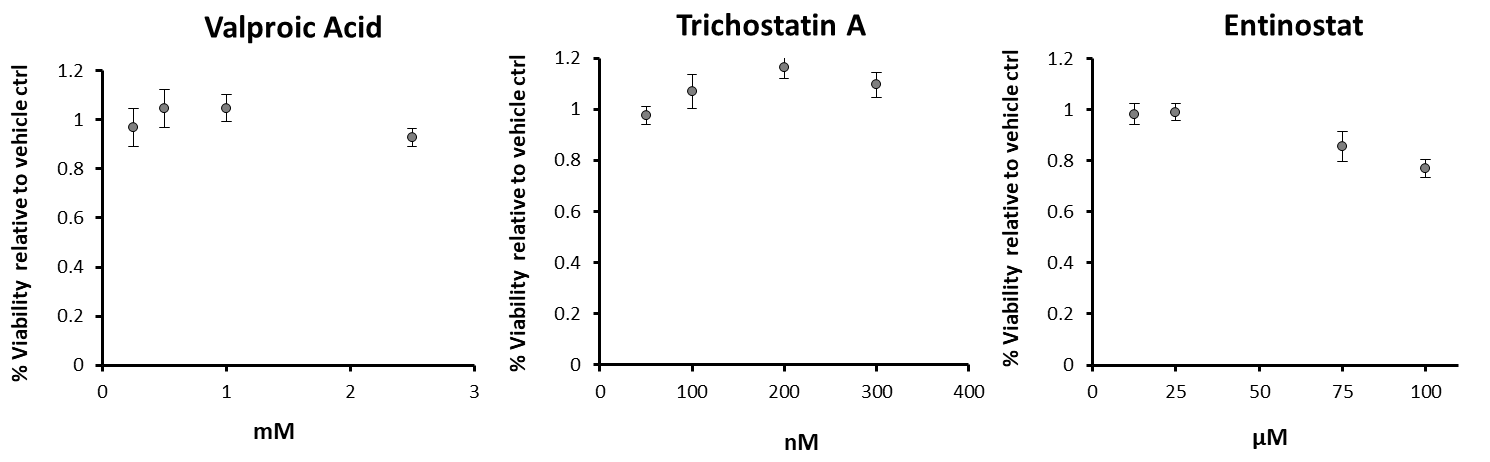

**Supplementary Figure 1**. Viability of TK6 cells after 4 hours exposures to valproic acid (left), trichostatin A (middle), and entinostat (right) followed by a 20-hour recovery period in fresh media, measured by the MTT assay (n = 6). Error bars represent standard deviation.
